# 6-O-galloylsalidroside, an active ingredient from Acer tegmentosum, ameliorates alcoholic steatosis and liver injury in a mouse model of chronic ethanol consumption

**DOI:** 10.1101/821397

**Authors:** Young Han Kim, Dong-Cheol Woo, Moonjin Ra, Sangmi Jung, Su Jung Ham, Ki Hyun Kim, Yongjun Lee

**Affiliations:** Hongcheon Institute of Medicinal Herb, 101 Yeonbongri, Hongcheon 25142, Republic of Korea; Convergence Medicine Research Center, Asan Institute for Life Sciences, Asan Medical Center, and Department of Convergence Medicine, University of Ulsan College of Medicine, Asan Medical Center, Seoul, Republic of Korea; Center for Bio-Imaging of New Drug Development, Asan Institute for Life Sciences, Asan Medical Centre, Seoul, Republic of Korea; School of Pharmacy, Sungkyunkwan University, Suwon 440-746, Korea

## Abstract

We previously reported that Acer tegmentosum extract, which is traditionally used to treat liver disease in Korea, may help reduce fat accumulation, improve liver metabolism, and suppress inflammation in alcoholic liver disease. The active ingredient was found to be 6-O-galloylsalidroside, which was isolated from the methanol extract of A. tegmentosum. We hypothesized that 6-O-galloylsalidroside extracted from A. tegmentosum may help protect from liver damage and attenuate hepatic fat accumulation associated with chronic alcohol consumption. In the present study, we aimed to investigate whether 6-O-galloylsalidroside can regulate alcoholic fatty liver and liver injury in mice. For this purpose, mice were fed with Lieber-DeCarli 5% ethanol diet for 11 days to induce steatosis and liver injury. Oral 6-O-galloylsalidroside was administered once a day for 11 days. Intrahepatic lipid accumulation caused by alcohol consumption was measured using in vivo 1H magnetic resonance imaging. Hepatic steatosis was observed histologically in tissue samples stained with hematoxylin and eosin, as well as Oil Red O. The serum levels of alanine aminotransferase (ALT) and aspartate aminotransferase (AST) were measured, as well as the triglyceride content in liver homogenates. On magnetic resonance spectroscopy, 6-O-galloylsalidroside appeared to alleviate alcohol-induced steatosis, which was reflected in decreased hepatic and serum triglyceride levels despite ethanol feeding. Furthermore, 6-O-galloylsalidroside treatment was associated with decreased RNA expression of Cd36, which plays an important role in the development of alcoholic steatosis through the hepatic de novo lipogenesis pathway. Furthermore, treatment with 6-O-galloylsalidroside inhibited the expression of cytochrome P4502E1 and attenuated hepatocellular damage, reflected in reduced ALT and AST levels. These findings suggest that 6-O-galloylsalidroside extracted from A. tegmentosum might serve as a bioactive agent for treating alcoholic fatty liver and liver damage.

## Introduction

Alcoholic hepatitis is a syndrome of progressive inflammatory liver injury associated with long-term heavy intake of ethanol [1-5]. Although genetic, environmental, nutritional, metabolic, and immunologic factors, as well as cytokines and viral diseases have been known to modulate the association between alcohol and liver disease, the exact mechanism underlying the development of alcoholic liver disease remains under debate [6-9]. The initial stage of alcohol-related diseases involves the development of fatty liver (steatosis) and accumulation of triglycerides in hepatocytes, which is a reversible condition. If alcohol consumption is continued, steatosis can progress to a potentially pathological condition such as steatohepatitis, fibrosis, cirrhosis, and even hepatocellular carcinoma, especially in the presence of other comorbidities [10-12].

Our previous findings indicated that Acer tegmentosum extracts protect against liver injury caused by alcohol consumption in mice [13]. In the process of isolation following extraction, 6-O-galloylsalidroside was purified and identified as the compound potentially responsible for the beneficial effect of A. tegmentosum extracts. The compound 6-O-galloylsalidroside is a phenolic glycoside found in several plants and has been studied for its potential antiviral and antioxidative activity [14-17]. Here, we hypothesized that 6-O-galloylsalidroside may serve as an active pharmaceutical ingredient to attenuate alcoholic steatosis and liver injury in vivo.

To test this hypothesis, we first established a chronic-plus-binge ethanol feeding model in mouse, characterized by alcohol-induced fat accumulation and immune responses underlying alcoholic liver injury, which leads to alcoholic hepatitis [18-20]. Furthermore, as the usefulness of in vivo 1H magnetic resonance imaging (1H-MRI) as a nondestructive and noninvasive analytical technique to measure fat levels in whole-liver tissue is well recognized (Garbow et al., 2004), we employed in vivo 1H-MRI to quantify alcohol-induced fat accumulation in the liver and thus identify a potential effect of 6-O-galloylsalidroside treatment in regulating hepatic lipid accumulation. Triglyceride levels in the serum and liver, as well as other relevant serum chemical parameters were analyzed to determine a potential effect of 6-O-galloylsalidroside treatment on these metabolic indicators.

Several groups have observed increased hepatic Cd36 expression at the transcriptional and translational levels in alcohol-fed mice. Cd36 is a class B scavenger receptor that may contribute to the lipid metabolism. It is thought that increased Cd36 expression promotes alcoholic steatosis through triglyceride synthesis and de novo lipogenesis in chronic alcohol-fed mice [21,22]. Therefore, we tested whether any effect of 6-O-galloylsalidroside on fat accumulation is mediated by suppression of Cd36 expression in chronic alcohol-fed mice.

Our present findings in a mouse model of chronic ethanol consumption indicate that 6-O-galloylsalidroside treatment protects against the development of alcoholic steatosis and liver damage, and that this effect is likely mediated by suppression of Cd36 RNA expression in the liver.

## Materials and Methods

### Isolation, Fractionation and Identification of 6-O-galloylsalidroside

The bark of A. tegmentosum was collected from Jeongseon in the Gangwon province, Korea in April 2013. The barks of A. tegmentosum (200 g) were dried at 60 °C for 24 h. The material was pulverized and extracted with distilled water (1 L) at 90 °C for 10 h and then filtered. The filtrate was concentrated in vacuo to afford resultant extracts (13.0 g). The resultant extracts (460 g) were suspended in distilled water (700 mL × 3) and successively solvent-partitioned with EtOAc, affording 22.4 g fractions. The EtOAc-soluble fraction was loaded to a Diaion HP-20 column chromatography, and fractionated with 0.5L each solvent system of 20%, 40%, 60%, 80%, and 100% MeOH in distilled water. Based on the results of a TLC analysis, the 80% and 100% MeOH miscible fractions were combined into one fraction. The combined fraction (4.2 g) was separated by RP-C18 silica gel (230–400 mesh) column chromatography eluting a gradient solvent system of MeOH-H2O (1:1–1:0, v/v) to afford three fractions (MeOH30, 70, 100 Fraction). MeOH70 Fraction (3.3 g) was subjected to silica gel (230–400 mesh) column chromatography and separated with a gradient solvent system of EtOAc-MeOH (30:1–1:1, v/v) to provide three fractions (A–C). Fraction C (928 mg) was passed through Sephadex LH-20 column chromatography using 100% MeOH to give eight subfractions (C1-8). 6-O-galloylsalidroside (7.7 mg, tR = 45.0 min) were purified from subfraction C5 (40 mg) by semipreparative reversed-phase HPLC (Phenomenex Luna Phenyl-hexyl, 250 × 10.0 mm, 5 μm) eluted with 38% MeOH/H2O (flow rate: 2 mL/min).

NMR spectra were recorded on a Varian UNITY INOVA 700 NMR spectrometer operating at 700 MHz (1H) and 175 MHz (13C), with chemical shifts given in ppm (δ). Preparative high performance liquid chromatography (HPLC) using a Waters 1525 Binary HPLC pump with a Waters 996 Photodiode Array Detector (Waters Corporation, Milford, CT, USA) was also performed. Semi-preparative HPLC was conducted using a Shimadzu Prominence HPLC System with SPD-20A/20AV Series Prominence HPLC UV-Vis Detectors (Shimadzu, Tokyo, Japan). Silica gel 60 (Merck, 70±230 mesh and 230±400 mesh) and RP-C18 silica gel (Merck, 40±63 μm) were used for column chromatography. The packing material for molecular sieve column chromatography was Sephadex LH-20 (Pharmacia, Uppsala, Sweden).

### Individual housing condition and experimental design

The experiment was performed in accordance with the ARRIVE guidelines proscribed by the Hongcheon Institute of Medicinal Herb Institutional Animal Care and Use Committee (approval no. HIMH 15-05). Male C57BL/6 mice were obtained from the Dae Han Biolink (Chungbuk, Korea). All mice were individually caged at controlled temperatures (23±1°C) with a 12-h light/dark cycle allowed to adapt to their new environment for one week. All mice were randomly divided into the following three groups (n = 7 or 8/group): Pair-fed group, Ethanol-fed group (5% vol/vol ethanol), Ethanol-fed + GAL group. GAL (10 mg/kg dose dissolved in distilled H2O) or vehicle was daily administered by oral gavage for 10 days.

### Alcoholic steatohepatitis and liver injury model

Male C57BL/6 mice, weighing 20–22 g were initially fed the control Lieber-DeCarli diet (Bio-Serv, Frenchtown, NJ, USA) ad libitum for 5 days to acclimatize to a liquid diet; subsequently, the mice were feeding ad libitum with Lieber-DeCarli ethanol diet containing 5% (vol/vol) or pair-fed a Lieber-DeCarli control diet for 10 days. On the morning of 11th day, mice were gavaged with a single dose of ethanol (5 g/kg body weight) or maltodextrin, respectively, and were euthanized 9 h later.

### Body weight and food intake evaluation

The body weight and food intake were measured daily between 4 PM and 6 PM. The food intake was calculated based on the measurements on the tube feeding (Bio-Serv, Frenchtown, NJ, USA), which was specifically designed to dispense liquid diets.

### Evaluation of serum and liver samples

On day 11 of the experiment, the animals were anesthetized using isoflurane and 500 µL of blood were collected from the orbital sinus. The blood was allowed to clot and then centrifuged at 4000 ×g for 10 min. The serum was separated and used for alanine transferase (ALT), aspartate transferase (AST), triglyceride(TG), and total cholesterol(TCHOL) assays using the Kornelab 20XT system (Thermo Fisher Scientific, Cleveland, OH, USA). Liver extracts were prepared by homogenization in 0.25% sucrose with 1 mmol/L EDTA and extracted as previously described (Miura et al., 2010). Hepatocyte triglyceride accumulation was quantified by extraction of hepatocyte lipids using chloroform/methanol (2:1). Liver of triglycerides was measured with the use of Triglyceride Determination Kit (Pointe Scientific, Canton, MI).

### RNA isolation and real-time RT-PCR

For real-time PCR analysis, equal aliquots of total RNA made from each mouse liver were pooled (total, 10 ug), using the TRIzol reagent (Invitrogen, Carlsbad, CA, USA), as recommended by the manufacturer. Total RNA was then reverse-transcribed and analyzed using real-time PCR with a Dice TP 800 Thermal Cycler and SYBR Premix ExTaq (Takara Bio, Osaka, Japan). The mRNA expression levels were calculated using *β* -actin as the control. The primer sequences used were as follows: for mouse Fas, forward 5′ -GGAGGTGGTGATAGCCGGTAT-3′ and reverse 5′ -TGGGTAATCCATAGAGCCCAG-3′; for mouse ApoB, forward 5′ -CGTGGGCTCCAGCATTCTA-3′ and reverse 5′ -TCACCAGTCATTTCTGCCTT-3′; for mouse Cd36, forward 5′ -TGGAGCTGTTATTGGTGCAG-3′ and reverse 5′ -TGGGTTTTGCACATCAAAGA-3′; and for mouse Fabp, forward 5′ -GCTGCGGCTGCTGTATGA-3′ and reverse 5′ -CACCGGCCTTCTCCATGA-3′.

### H&E, Cyp2e1 and Oil Red O staining

Liver organs were fixed with 10% neutral-buffered formalin for 24 h prior to paraffin embedding using standard procedures. Liver sections (4 µm in thickness) were stained with hematoxylin and eosin. Cyp2e1 staining was performed by using anti-Cyp2e1 antibody, and incubated with an appropriate biotin-conjugated rabbit anti-goat IgG antibody (Burlingame, CA, USA), indicated in the manufacturer’s procesures. For Oil Red O staining, liver tissues were frozen in OCT compounds, and cut at 5 µm in thickness. The sections were mounted on slides drying for 2 hrs and fixed with 10% formalin. After fixation, the slides were stained with 0.5% Oil Red O solution (Sigma-Aldrich, St. Louis, MO, USA) for 20 min, and then counterstained with hematoxylin (DAKO, Carpinteria, CA, USA).

### ^1^H magnetic resonance imaging and spectroscopy

During magnetic resonance scanning, all mice were anesthetized by inhalation of 1– 1.5% isoflurane mixed in air and delivered *via* a mask. The animals breathed freely and were monitored for changes in the respiratory rate to adjust the concentration of anesthetic in the magnetic resonance machine. All scans were performed in a horizontal, 9.4-T, 160-mm, magnetic resonance system for small animals (Agilent, Palo Alto, CA, USA). For MRI-based estimation of proton density fat fraction, fast spin-echo images were acquired using the following settings: repetition time (TR), 3000 ms; echo time (TE), 10.5 ms; matrix size, 192×192; 24 segments; echo train length, 8; slice thickness, 2.0 mm; number of excitations, 4; 6 slices; field of view, 20 mm × 30 mm. Fat signal was suppressed using an offset sinc pulse (duration, 4.0 ms) at −1450 Hz. Two images (fat suppressed/unsuppressed) were acquired for each position in order to calculate the fat fraction factor.

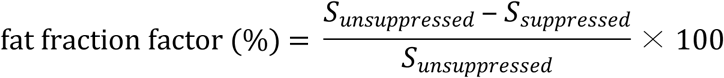

where *S*_*unsuppressed*_ and *S*_*suppressed*_ represent, respectively, the fat unsuppressed and suppressed magnetic resonance signal intensity.

### Statistical Analysis

All statistical analyses were performed using GraphPad Prism version 5.0 (GraphPad Software, San Diego, CA, USA). The results were expressed as the mean ± standard deviation. Comparisons between groups were performed using the unpaired Student t-test. One-way analysis of variance (ANOVA) was used for comparisons between multiple groups. Statistical significance was established at p < 0.05. All data appeared to be normally distributed. Each variable of interest was analyzed using the most appropriate statistical test.

## Results

### Isolation of structural identification of 6-O-galloylsalidroside

Dried and pulverized A. tegmentosum bark was extracted with water at 90 °C and then filtered. The filtrate was concentrated under vacuum to give crude aqueous extract and then solvent-partitioned with hexane, CH2Cl2, EtOAc, and n-BuOH, affording four main fractions. According to preliminary 1H NMR spectroscopic analysis, the EtOAc-soluble fraction was selected to identify bioactive constituents. Fractionation and purification on repeated column chromatography and semi-preparative HPLC of the EtOAc fraction resulted to the isolation of A-C fractions (Fig. 1A). The isolated compounds were identified as 6-O-galloylsalidroside[17] by comparing their NMR spectroscopic and physical data with those in the literature and measurement of their specific rotations (Fig. 1B).

**Fig 1.**
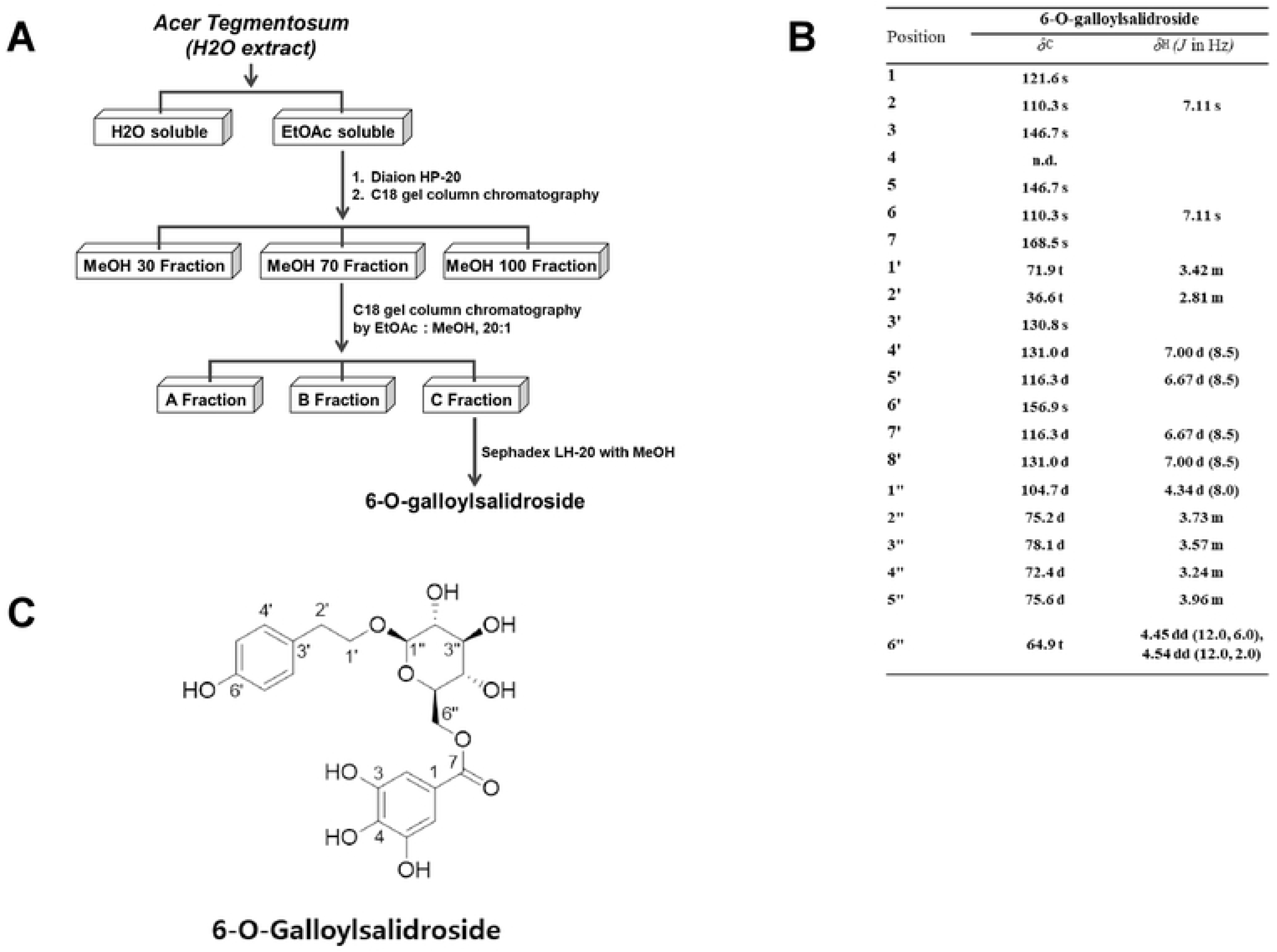
Active compound isolation and identification. (A) A scheme of the chromatographic methods used for the separation of *A. tegmentosum*’s active compound. (B) ^1^H(700MHz) and ^13^C(175MHz) NMR data of 6-O-galloylsalidroside in CD_3_OD.

### Beneficial effect of 6-O-galloylsalidroside on hepatic fat accumulation and metabolic indicators e

All ethanol-fed mice, including those treated with 6-O-galloylsalidroside, had a slightly lower body weight than noted among pair-fed controls; however, there was no difference between Ethanol-fed + GAL group and Ethanol-fed group (Fig. 2A). Food intake was the same in all groups. Furthermore, there were no significant between-group differences regarding the liver, kidney, or spleen relative weight (Fig. 2C). We conducted 1H-MRI-based measurements of proton density fat fraction to confirm the establishment of alcoholic fatty liver. There was no between-group difference in fat liver volume at the beginning of the experiment (i.e., after 4 days during which all mice consumed control Lieber-DeCarli diet). However, at 11 days, the volume of hepatic fat was higher in ethanol-fed mice than in pair-fed controls (Fig. 3A), reflected in a three-fold higher fat fraction factor (Fig. 3B), whereas 6-O-galloylsalidroside-treated ethanol-fed mice had a fat fraction factor as low as that of pair-fed controls. The histopathology and Oil Red O staining findings were highly consistent with the 1H-MRI data (Fig. 3C).

**Fig 2.**
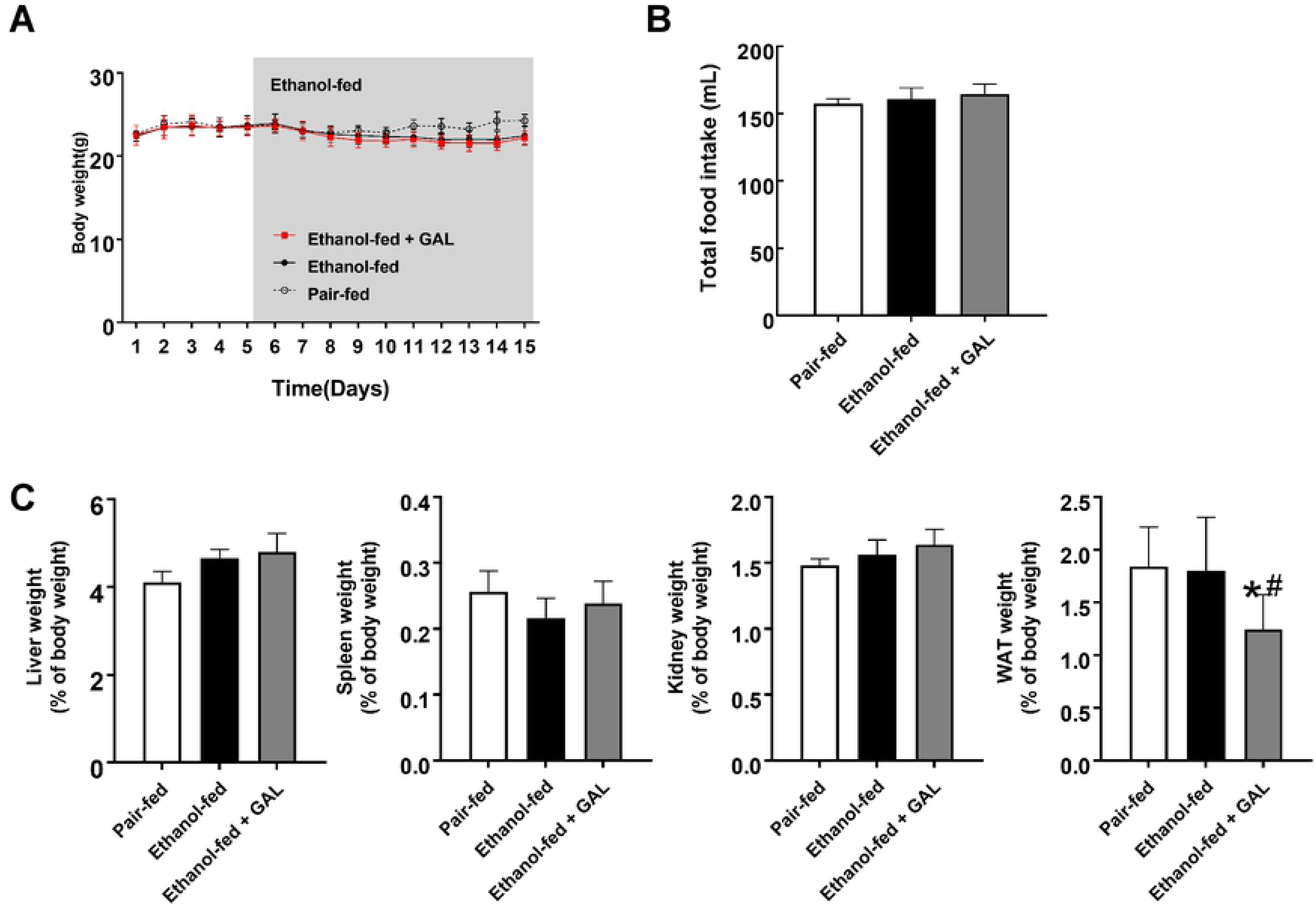
Treatment with 6-O-galloylsalidroside has no effect on body weight or food intake in ethanol-fed mice. (A) Effect of 6-O-galloylsalidroside on body weight. (B) Effect of 6-O-galloylsalidroside on food intake. (C) Effect of 6-O-galloylsalidroside on the liver, spleen, kidney, and white adipose tissue relative weight. Relative organ weight was obtained as the weight of the organ divided by the body weight and expressed as a percentage. Data are shown as mean ± standard deviation (n = 7). Statistical significance: ^#^p < 0.05 and *p < 0.05. Abbreviations: GAL, 6-O-galloylsalidroside; WAT, white adipose tissue.

**Fig 3.**
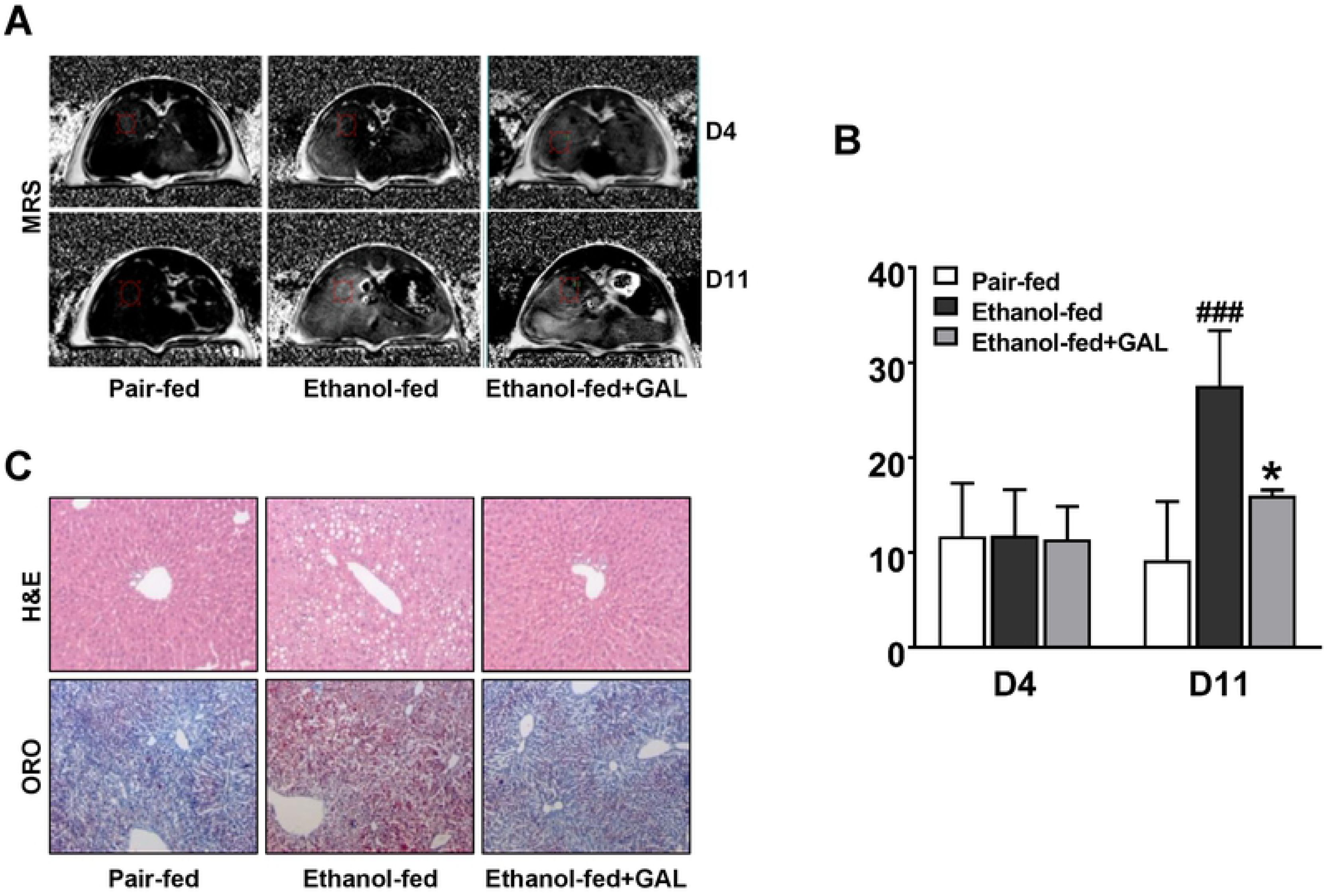
Fat deposition in the liver is attenuated by 6-O-galloylsalidroside treatment. (A) In vivo ^1^H MRI (proton density imaging) of mouse liver. (B) Fat fraction factors in the liver, based on FFPD. (C) Representative photomicrographs of stained liver sections. Error bars represent the standard deviation with each group. Statistical significance: ^###^p < 0.001, *p < 0.05. Abbreviations: GAL, 6-O-galloylsalidroside; H&E, hematoxylin and eosin; MRS, magnetic resonance spectroscopy; ORO, Oil Red O.

Among ethanol-fed mice, significantly lower levels of both serum and hepatic triglycerides were detected in Ethanol-fed + GAL group than in Ethanol-fed group (Fig. 4A, B). However, no significant between-group differences were observed regarding total cholesterol levels in the serum (Fig 4B).

**Fig 4.**
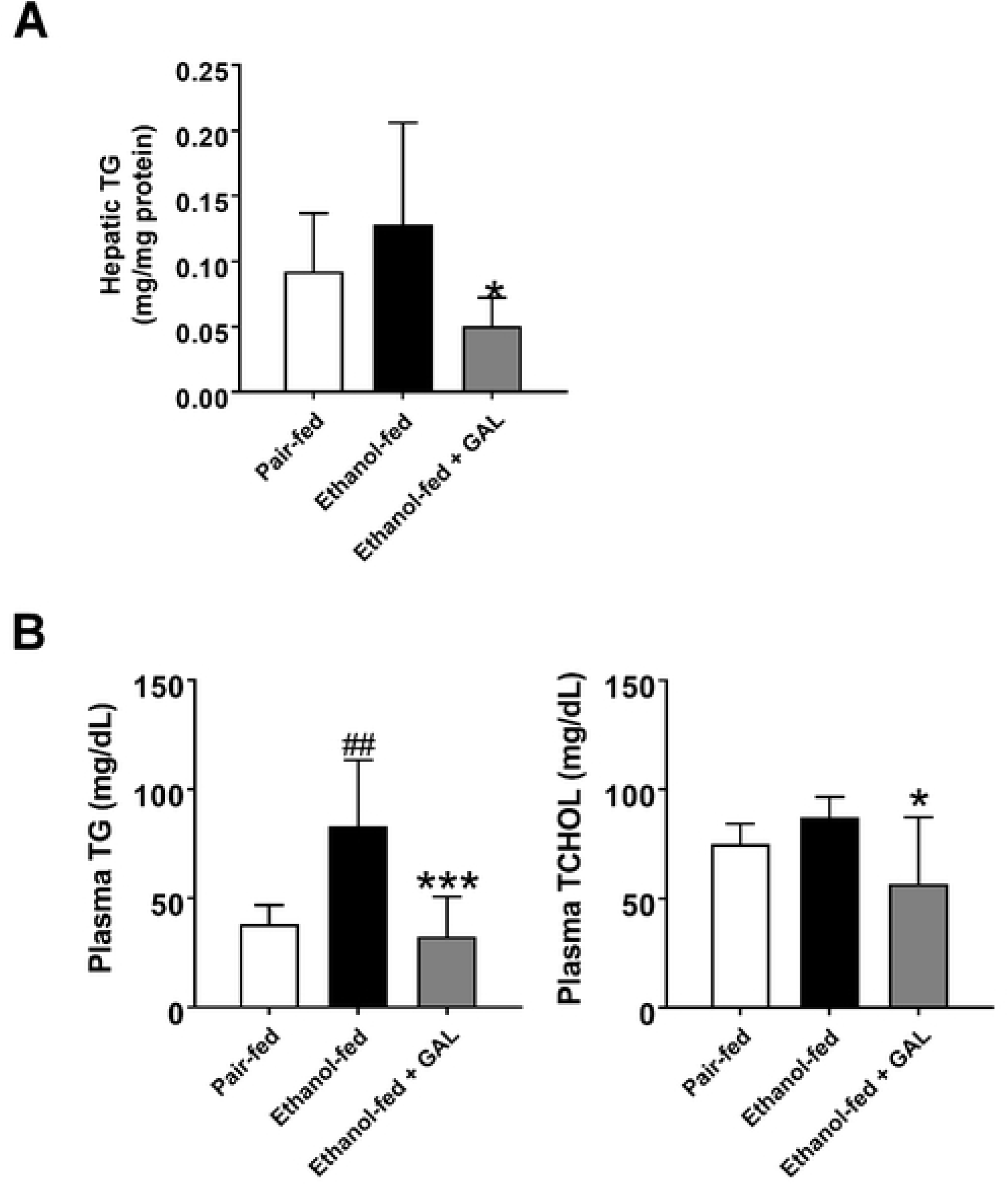
Triglyceride levels in serum and liver are reduced by 6-O-galloylsalidroside treatment. (A) Effect of 6-O-galloylsalidroside on hepatic triglyceride levels. (B) Effect of 6-O-galloylsalidroside on serum triglyceride and total cholesterol levels. Data are shown as mean ± standard deviation (n = 7). Statistically significant differences versus Pair-fed group and Ethanol-fed group are marked as ^##^p < 0.01, *p < 0.05 and ***p < 0.001, respectively. Abbreviations: GAL, 6-O-galloylsalidroside; TCHOL, total cholesterol; TG, triglycerides.

### Alcohol-induced elevation of liver Cd36 gene expression is suppressed by 6-O-galloylsalidroside

Our data indicated that 6-O-galloylsalidroside affects liver fat deposition associated with chronic alcohol consumption. We then performed RT-PCR to determine whether this effect of 6-O-galloylsalidroside was reflected in the expression of genes involved in alcohol and fat metabolism in the liver.

We found that the gene expression of Cd36 was significantly higher in Ethanol-fed group than in Pair-fed group, but this effect was significantly reduced in Ethanol-fed + GAL group (by 57%; p < 0.05). Taken together with other recent findings, the present observations indicate that 6-O-galloylsalidroside may be an important regulator of alcoholic steatosis by driving de novo lipogenesis mediated by Cd36. Abbreviations: GAL, 6-O-galloylsalidroside (Fig 5).

**Fig 5.**
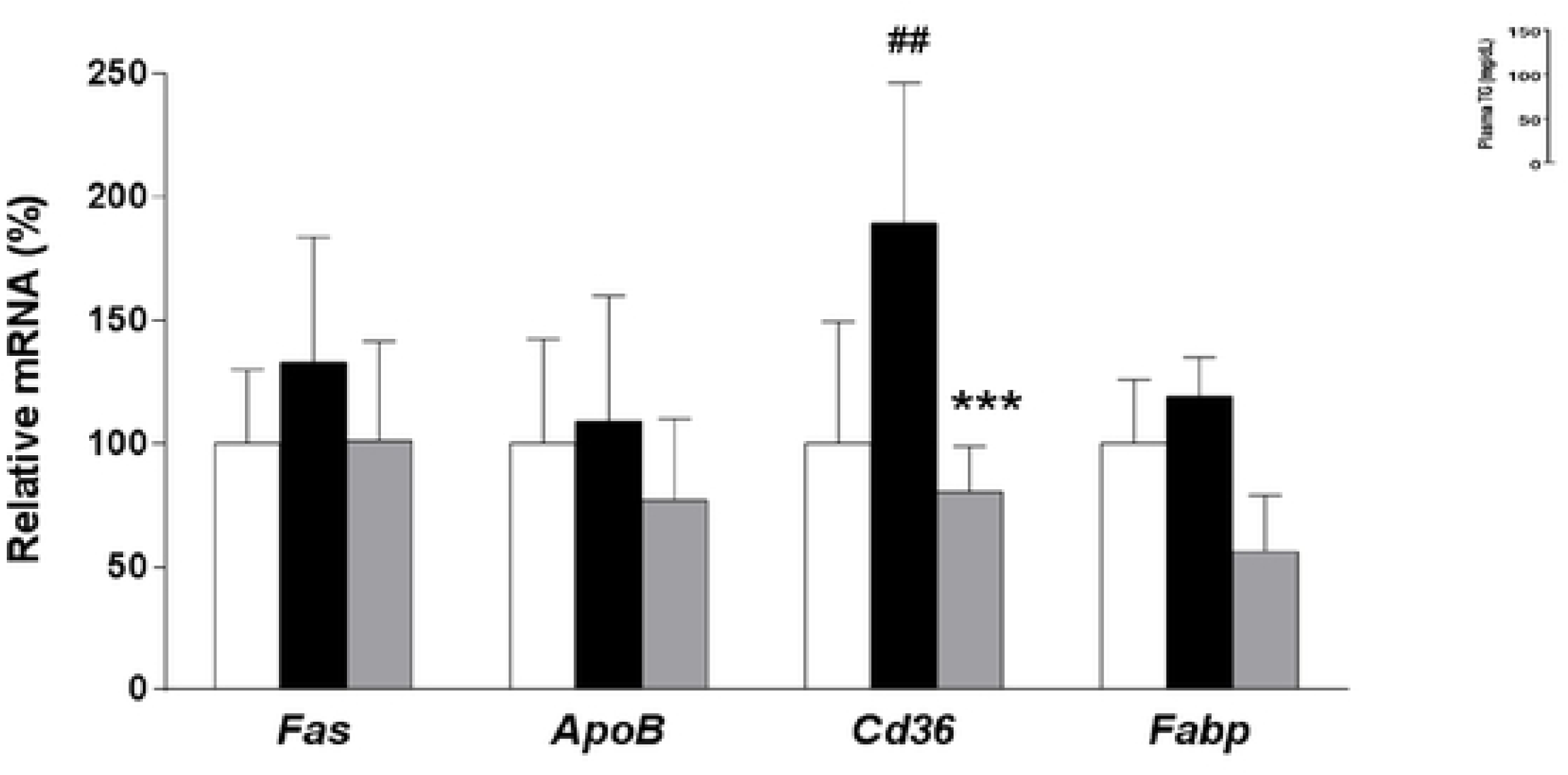
Treatment with 6-O-galloylsalidroside suppresses the increase in *Cd36* gene expression induced by chronic alcohol consumption. Quantitative real-time PCR analysis of genes involved in alcohol and lipid metabolism. Data are shown as mean ± standard deviation (n = 7). Statistically significant differences versus Pair-fed group and Ethanol-fed group are marked as ^##^p < 0.01 and ***p < 0.001, respectively. Abbreviations: GAL, 6-O-galloylsalidroside.

### Protective effect of 6-O-galloylsalidroside against alcohol-induced liver injury

We investigated whether 6-O-galloylsalidroside affected the serum levels of ALT and AST.

The levels of ALT and AST were significantly higher in Ethanol-fed group than in Pair-fed group. This effect was reduced in ethanol-fed mice treated with 6-O-galloylsalidroside. Although the reduction was not statistically significant, the levels of ALT and AST in ethanol-fed mice treated with 6-O-galloylsalidroside were similar to those in pair-fed controls (Fig. 6A). Immunohistochemistry evaluation revealed no differences in Cyp2e1 expression between Ethanol-fed + GAL group and Ethanol-fed group (Fig. 6B).

**Fig 6.**
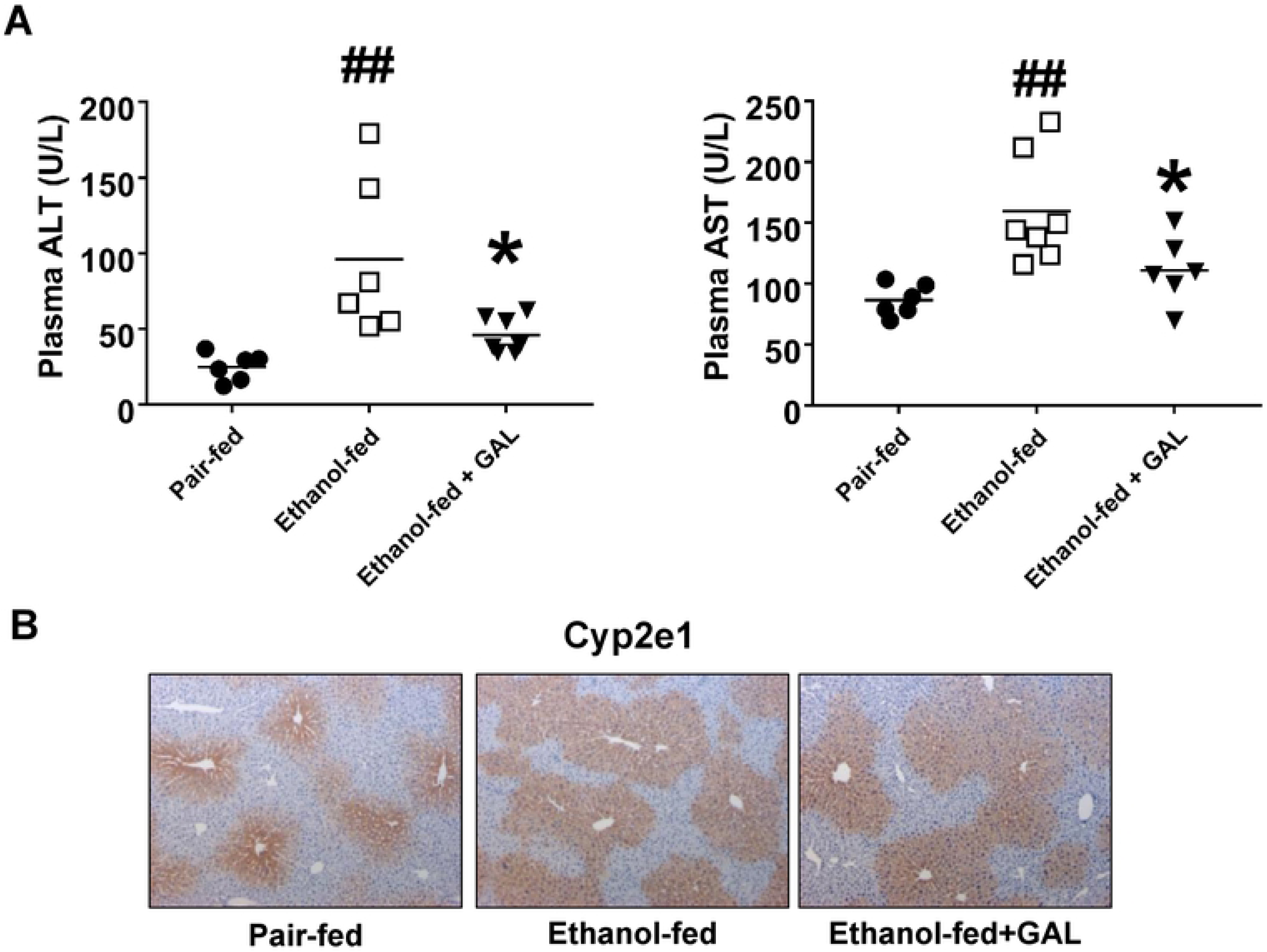
Treatment with 6-O-galloylsalidroside attenuates alcohol-induced liver damage. (A) Effects of 6-O-galloylsalidroside treatment on the serum levels of ALT and AST. (B) Effect of 6-O-galloylsalidroside on Cyp2e1 expression in liver tissue, as determined by immunohistochemistry. Original magnification, ×100. Data are shown as mean ± standard deviation (n = 8). Statistically significant differences versus Pair-fed group and Ethanol-fed group are marked as ^##^p < 0.01 and *p < 0.05, respectively. Abbreviations: ALT, alanine aminotransferase; AST, aspartate aminotransferase; GAL, 6-O-galloylsalidroside.

## Discussion

6-O-galloylsalidroside is a phenolic compound isolated from the dried twigs of A. tegmentosum [23]. 6-O-galloylsalidroside leads to radical scavenging activities and reducing inflammation in under the inflammatory condition [24]. The true prevalence of alcoholic hepatitis worldwide remains unknown because patients may be asymptomatic and/or may never seek medical attention. Therefore, only patients with severe alcoholic hepatitis are likely to be prescribed steroid drugs such as prednisolone to reduce liver injury and suppress inflammation. However, clinical studies have been reported serious adverse effects like an increased susceptibility to infection in patients with alcoholic hepatitis. Our previous findings, as well as several other studies [25] [17] [13] [26] [23], indicated that A. tegmentosum extract may be useful to treat hepatic disorders such as hepatic inflammation, hepatic cirrhosis, and hepatic cancer. Indeed, we suggested that quercetin-3-O-β-D-xylopyranoside isolated and identified from A. tegmentosum, may be an active compound that ameliorates alcohol-induced fatty liver and liver injury by improving hepatic energy metabolism and preventing hepatic apoptosis.

6-O-galloylsalidroside has been previously patented as a pharmaceutical composition for preventing or treating liver disease. 6-O-galloylsalidroside prevented hepatic TG synthesis and liver injury during chronic alcohol consumption.

In this study, we conducted 1H-MRI-based measurements of proton density fat fraction to confirm the establishment of alcoholic fatty liver via chronic-plus-binge alcohol feeding, as well as to determine whether 6-O-galloylsalidroside treatment caused a reduction in hepatic fat accumulation in ethanol-fed mice [27]. 6-O-galloylsalidroside, used at the same dosage used in the patent, led to complete reduction of hepatic fat accumulation by chronic alcohol consumption. We found that Cd36 expression was higher in alcohol-fed mice than in pair-fed controls, whereas Ethanol-fed + GAL group had relatively normal Cd36 expression despite chronic alcohol consumption. Finally, upon examining the biomarkers of alcohol-induced liver injury, we concluded that 6-O-galloylsalidroside treatment protects from liver apoptosis caused by excessive alcohol consumption, by inhibiting the alcohol-induced increase in ALT and Cyp2e1 enzymes [22].

It should be noted that, when judging the efficacy of 6-O-galloylsalidroside treatment in alcoholic fatty liver, we only evaluated the RNA expression levels of the Cd36 gene and the amount of hepatic fat accumulation. Thus, although our results are very promising, they may not necessarily indicate that 6-O-galloylsalidroside treatment regulates lipogenesis directly through the Cd36 pathway. Furthermore,

Taken together, our present results indicate that 6-O-galloylsalidroside extracted from A. tegmentosum inhibits hepatic lipogenesis as well as liver inflammation caused by alcohol consumption, thus protecting the liver from alcohol-induced apoptosis and injury. Our study was the first to demonstrate that 6-O-galloylsalidroside ameliorates alcoholic fatty liver and attenuates liver injury by preventing de novo lipogenesis and apoptosis in the liver, highlighting the possible use of 6-O-galloylsalidroside as an active pharmaceutical ingredient for the prevention and treatment of alcohol-related diseases.

